# The Ndc80 complex targets Bod1 to human mitotic kinetochores

**DOI:** 10.1101/131664

**Authors:** Katharina Schleicher, Sara ten Have, Iain M. Porter, Jason R. Swedlow

## Abstract

Regulation of protein phosphatase activity by endogenous protein inhibitors is an important mechanism to control protein phosphorylation in cells. We recently identified Biorientation defective 1 (Bod1) as a small protein inhibitor of PP2A-B56. This phosphatase controls the amount of phosphorylation of mitotic kinetochore proteins and thus the establishment of load-bearing chromosome-spindle attachments in time for accurate separation of sister chromatids in mitosis. The PP2A-B56 regulatory protein Bod1 directly localises to mitotic kinetochores and is required for correct segregation of mitotic chromosomes. In this report, we have probed the spatio-temporal regulation of Bod1 during mitotic progression. Kinetochore localisation of Bod1 increases from nuclear envelope breakdown until metaphase. Phosphorylation of Bod1 at threonine 95 (T95), which increases Bod1’s binding and inhibition of PP2A-B56, peaks in prometaphase, when PP2A-B56 localisation to kinetochores is highest. Kinetochore targeting of Bod1 depends on the outer kinetochore protein Ndc80 and not PP2A-B56. Additionally, Bod1 depletion leads to a defect in the phosphorylation kinetics of Ndc80 at serine 55 (S55) within its N-terminus. Therefore, Ndc80 recruits a phosphatase inhibitor to kinetochores which directly feeds forward to regulate Ndc80 phosphorylation and the stability of the attachments of microtubules to kinetochores.

## 1. Introduction

To preserve genome integrity, the two sister chromatids of each mitotic chromosome must be distributed equally between daughter cells. Movement of sister chromatids to opposite poles of a dividing cell requires attachment to spindle microtubules of opposing orientation. Errors in the attachment process can lead to chromosome missegregation and aberrant chromosome numbers. Such aneuploid karyotypes are the major cause of spontaneous miscarriages in humans [1] and often observed in cancer genomes [2]. Spindle microtubule attachment occurs at a multi-complex protein interface between mitotic chromosomes and the spindle apparatus called the kinetochore [3]. The kinetochore consists of approximately 30 core structural proteins that are arranged into several functional subcomplexes. As well as providing a structural link between chromosomes and spindle microtubules, structural kinetochore proteins also act as a signalling platform by recruiting checkpoint proteins, kinases, and phosphatases. Early kinase activity destabilises kinetochore-microtubule interactions [4,5] to prevent attachment errors [6]. Conversely, phosphatase activity is needed at later stages of the attachment process to stabilise kinetochore-microtubule interactions that have passed the quality-control and thus protect load-bearing attachments [7]. There is a conserved, dynamic system comprised of at least eight kinases and two phosphatases that control microtubule-kinetochore attachments [3,8].

The detailed role and function of protein phosphatases and their interplay at the kinetochore is only beginning to be elucidated [9–11], but both are of great interest as they are absolutely required to ensure faithful chromosome segregation. Protein phosphatase 1 (PP1) dephosphorylates microtubule-binding kinetochore proteins to ultimately stabilise attachments [12,13]. However, recruitment of PP1 to kinetochores requires the initial activity of protein phosphatase 2A (PP2A) [14], highlighting the importance of coordinated timely activation of these kinetochore components. PP2A is a heterotrimeric enzyme, composed of a scaffolding (A) subunit, catalytic (C) subunit, and a regulatory (B) subunit [15]. It is targeted to the kinetochore by the B56 family of B subunits [11,16,17]. The highest mitotic occupancy of PP2A-B56 at kinetochores is reached in prometaphase and can be maximised by increasing the number of unattached kinetochores with nocodazole [11]. Under the same conditions, PP1 localisation to kinetochores is low [12], suggesting that PP2A accumulation at unattached kinetochores alone is not sufficient to recruit PP1 and that additional molecular signals are required to activate PP2A-mediated PP1 recruitment in metaphase.

We have recently identified a small kinetochore protein, Biorientation defective 1 (Bod1), that can specifically inhibit PP2A-B56 [9,18]. Bod1 is required for cognitive function in humans and *Drosophila* models [19] and is thought to contribute to the early stages of radiation-induced genomic instability [20]. Depletion of Bod1 from HeLa cells leads to premature loss of phosphorylation on several kinetochore proteins, including MCAK and CENPU, due to unregulated activity of PP2A-B56. It also causes an increase in aberrant chromosome attachments and defective chromosome segregation. Bod1, together with CIP2A [21], FAM122A [22], I1PP2A/ANP32A [23], I2PP2A/SET [24], TIP [25], and Arpp-19/Ensa [26,27], forms part of a growing family of PP2A inhibitors that have important roles in supporting cell division. However, little is known about the localisation of these PP2A regulators or how they modulate the activity of PP2A towards different substrates.

We have studied the temporal recruitment and phospho-regulation of Bod1 at mitotic kinetochores. Bod1 is recruited to kinetochores by the outer kinetochore protein Ndc80 (Nuclear division cycle protein 80, also known as highly expressed in cancer protein Hec1). Furthermore, Bod1 can protect phosphorylation of a key site in the N-terminal tail of Ndc80 that is required for microtubule attachment. These data refine our understanding of how PP2A activity at the kinetochore is regulated and identify another target of the Bod1 phosphatase inhibitor pathway.

## 2. Results

### Bod1 localises to kinetochores throughout mitosis and is maximally phosphorylated in prometaphase

To dissect the temporal regulation of Bod1 recruitment to kinetochores, we raised peptide antibodies for immunofluorescence profiling in HeLa cells (figure 1*a*; electronic supplementary material, figure S1). We quantified Bod1 kinetochore intensities within a 4 pixel radius of anti-centromere antibody (ACA) staining as a reference (figure 1*c*). Bod1 is first detected on kinetochores at nuclear envelope breakdown and reaches its maximum at metaphase.

**Figure 1.**
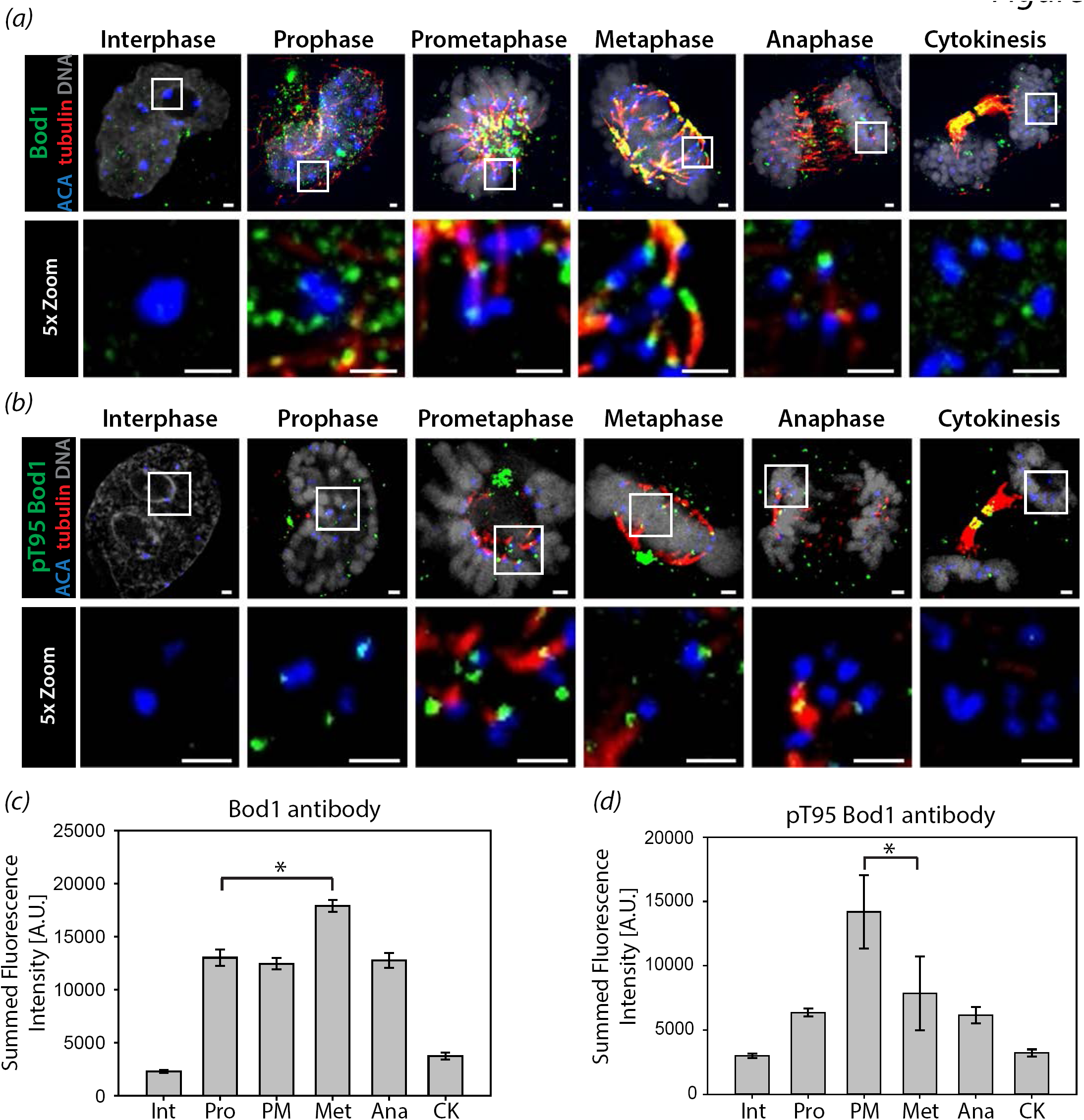
Cell cycle profiles of Bod1 kinetochore recruitment and phosphorylation. HeLa cells were fixed in paraformaldehyde and stained with (*a*) a Bod1 pan-specific peptide antibody raised in sheep or (*b*) a T95 phospho-specific antibody (both green). Mitotic cells were imaged and the kinetochore was marked as the interface between centromeric region (anti-centromere antibody, ACA, in blue) and spindle microtubules (tubulin, red). Top panel shows a single z-section of each cell cycle stage. Bottom panels are magnifications of the same cell (section indicated by white boxes). Scale bars are 1 μm. Quantification of (*c*) total Bod1 or (*d*) T95-specific phospho-Bod1fluorescence intensity at the kinetochore corresponding to experiments shown in (*a*) and (*b*). Background-corrected summed intensity of centromeric Bod1 staining was quantified. n = 10 cells per mitotic phase. Error bars represent standard error. Asterisks indicate significance in multiple comparison after ANOVA on ranks. Int: interphase, Pro: prophase, PM: prometaphase, Met: metaphase, Ana: anaphase, CK: cytokinesis.

We showed previously that inhibition of PP2A-B56 by Bod1 is greatly enhanced by its phosphorylation at T95 [9]. We therefore raised a phospho-specific antibody against this site (figure 1*b*; electronic supplementary material, figure S1). Quantification of pT95 Bod1 at kinetochores revealed that this post-translational modification peaks in prometaphase, before maximal recruitment of the total protein (figure 1*d*). PP2A-B56 levels at kinetochores are highest in prometaphase when attachments are weak [11]. The phosphorylation of Bod1 at T95 therefore coincides with the recruitment of PP2A-B56, consistent with a role in inhibiting PP2A-B56 activity and enabling correction of attachment errors in early mitosis.

### Bod1 recruitment to kinetochores is independent of PP2A-B56 and Knl1

To test whether Bod1 and PP2A-B56 are co-recruited to kinetochores, we depleted PP2A-B56 from HeLa cells using a pool of B56 isoform-specific siRNAs [11] and quantified total Bod1 protein at the kinetochores (figure 2*a-e*). Surprisingly, there was no significant change in Bod1 recruitment to kinetochores upon B56 depletion. Since it is difficult to achieve complete knockdown of B56 isoforms via siRNA (figure 2*e*), we then depleted the outer kinetochore protein Knl1, which acts as a scaffolding platform that recruits PP2A-B56 to kinetochores via BubR1 [16,17,28]. As with B56 depletion, siRNA-mediated knockdown of Knl1 did not affect Bod1 recruitment to kinetochores (figure 2*f-h*). We therefore conclude that Bod1 is recruited to kinetochores independently of its target, PP2A-B56, and via a different interaction platform.

**Figure 2.**
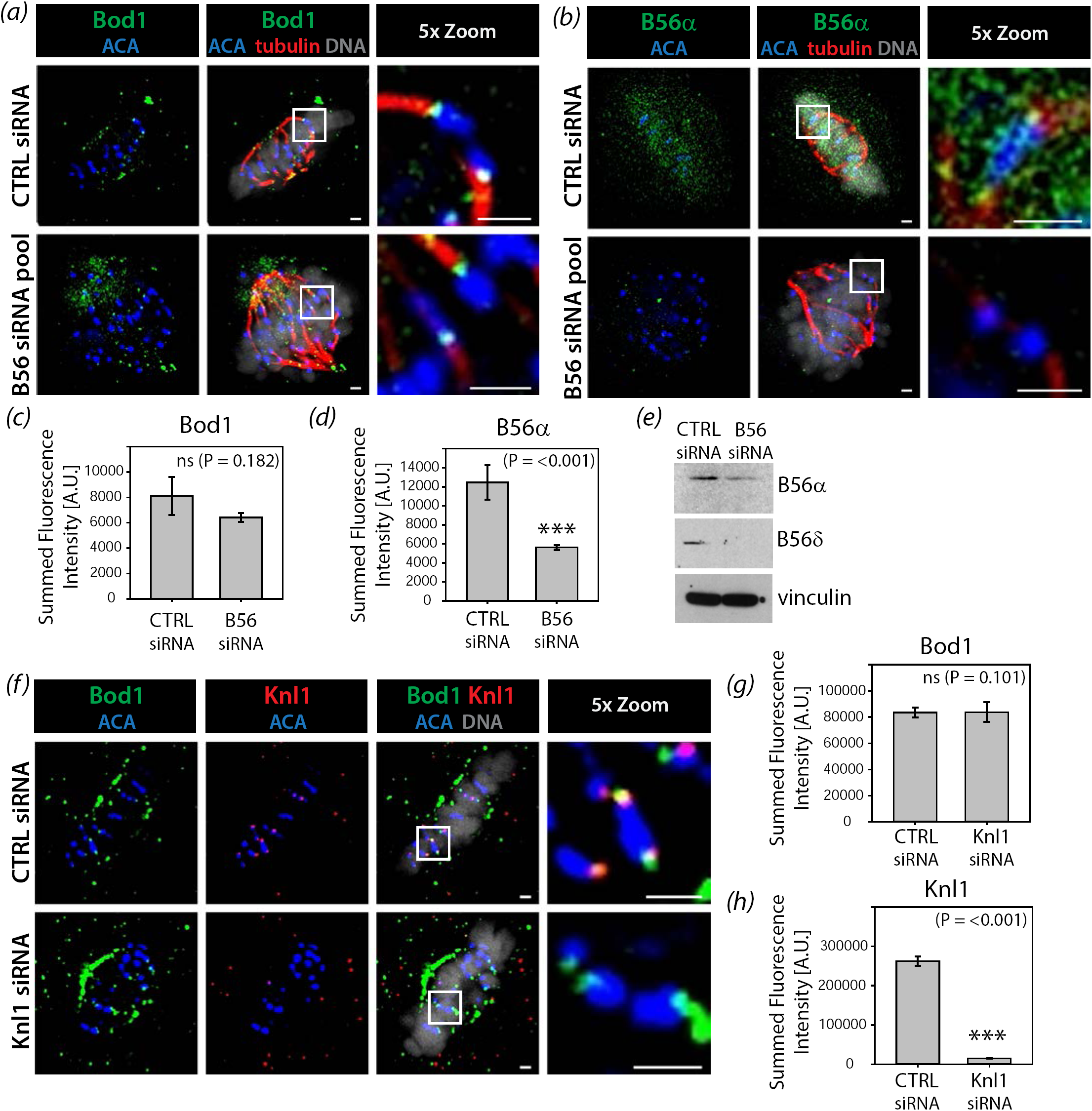
PP2A-B56 and Knl1 are dispensable for Bod1 recruitment to kinetochores. HeLa cells were treated with either control siRNA or a smart pool siRNA targeting all PP2A-B56 isoforms as described previously [11]. After 48h cells were fixed in paraformaldehyde and stained with (*a*) the total Bod1 peptide antibody or (*b*) a PP2A-B56α isoform-specific antibody. Metaphase control cells or B56siRNA-treated cells showing the characteristic B56-depletion phenotype of metaphase chromosome alignment defects were imaged. The last panels are magnifications of the same cell (section indicated by white boxes). Scale bars are 1 μm. Total (*c*) Bod1 and (*d*) B56α intensities at kinetochores were quantified. n = 10 cells per condition. Statistical test used was Student’s t-test. Three asterisks indicate highly significant difference. ns: no significant difference in fluorescence intensity. Error bars represent standard error. (*e*) Immunoblot corresponding to the B56 depletion experiments in (*a-d*) using antibodies against two of the five targeted B56 isoforms and vinculin as a loading control. (*f*) HeLa cells were treated with either control or Knl1 siRNA for 48h, fixed in paraformaldehyde, and co-stained with both total Bod1 peptide antibody and a Knl1-specific antibody. Metaphase cells were imaged for both treatment conditions. The last panels are magnifications of the same cell (section indicated by white boxes). Scale bars are 1 μm. Total (*g*) Bod1 and (*h*) Knl1 intensities at kinetochores were quantified. n = 10 cells per condition. Statistical test used was Student’s t-test. Three asterisks indicate highly significant difference. ns: no significant difference in fluorescence intensity. Error bars represent standard error.

### The mitotic interactome of Bod1 contains many outer kinetochore proteins including Ndc80

To discover candidate proteins that might target Bod1 to kinetochores, we combined affinity purification of Bod1 with label-free quantitative mass spectrometry (MS) (figure 3*a*; electronic supplementary material, figure S2). In mitotic lysates from HeLa cells expressing Bod1-GFP, we identified and could quantify 3512 proteins. Of these, 42 were significantly enriched in affinity purifications from Bod1-GFP expressing cells compared to cells expressing GFP alone as a control (n = 4 biological replicates; electronic supplementary material, table S1). Analysis of gene ontology (GO) term annotation identified 95 centromere- and kinetochore-associated proteins in the Bod1-GFP affinity purifications (electronic supplementary material, table S2). Of these, Bod1 itself, Nuclear division cycle protein 80 (Ndc80; also known as Highly expressed in cancer protein (Hec1)), and Dynein intermediate chain 1 (DYNC1I1) were significantly enriched in Bod1-GFP affinity purifications compared to controls (figure 3*b*; electronic supplementary material, figure S3*a*). The most reproducible kinetochore interactor was Ndc80; it was found in all four biological replicates of the experiment. Furthermore, of all kinetochore proteins detected, Ndc80 exhibited the highest fold change in Bod1-GFP affinity purifications compared to controls. Intensity analysis of the centromeric region in HeLa cells, co-stained with Bod1 and Ndc80 antibodies, revealed that the immunofluorescence signal of the two proteins overlaps at the outer kinetochore (figure 3*c*). The mitotic Bod1 interactome also contained components of the SET1B methyltransferase complex with significant enrichment of ASH2L. This is consistent with previous results in asynchronous HeLa cells [29].

**Figure 3.**
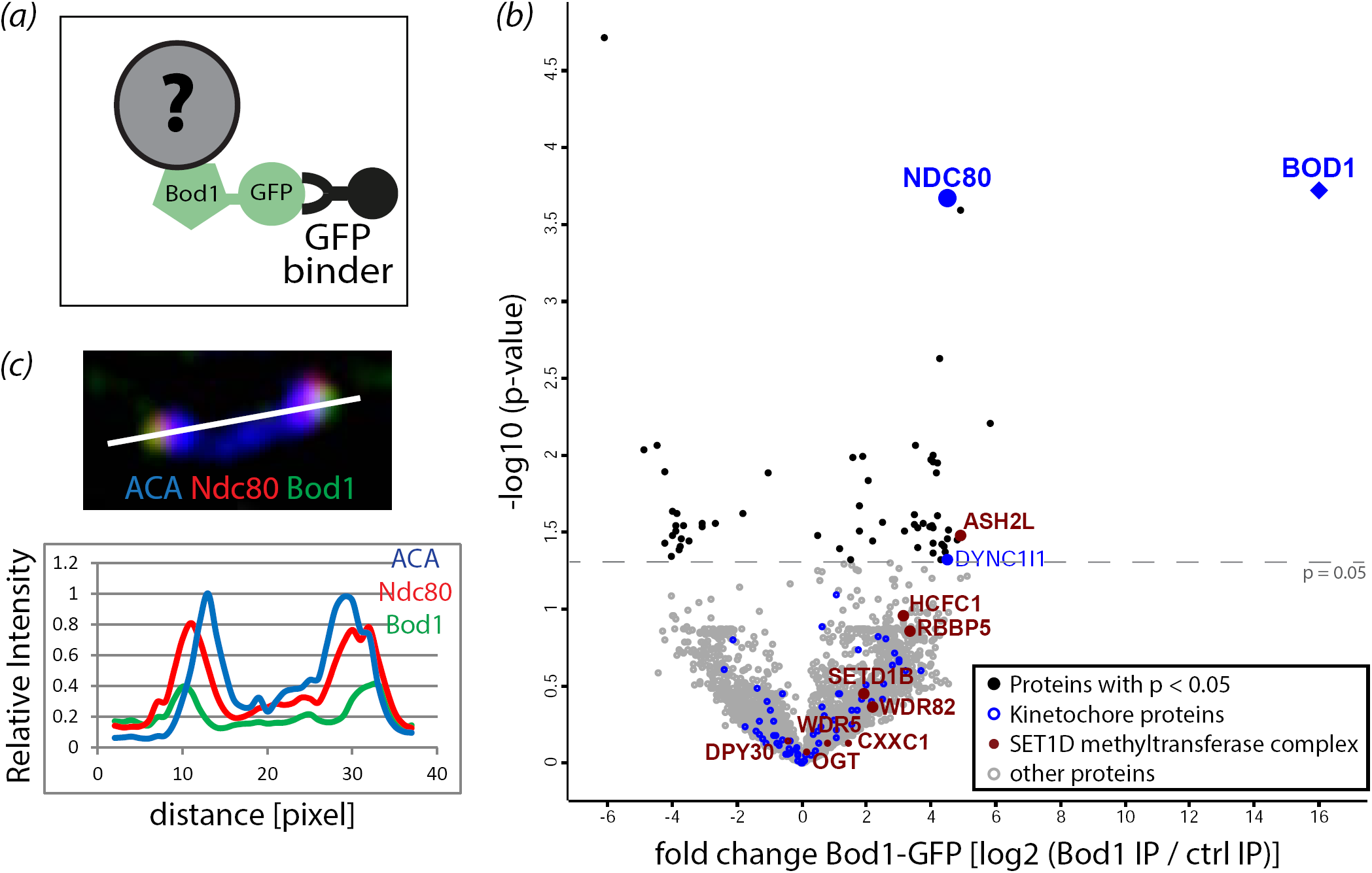
Proteomics analysis of mitotic Bod1-GFP affinity purifications. (*a*) GFP binder was used to affinity purify Bod1-GFP and associated proteins from mitotic HeLa cell lysates as described previously [9]. (*b*) Volcano plot of the mitotic Bod1 interactome detected by LFQ shotgun mass spectrometry. The fold change in Bod1-GFP samples vs control (represented by log_2_ of their intensity ratios) is plotted against –log_10_ of the p-value obtained in an unpaired Student’s t-test with a threshold p-value of 0.05. All kinetochore proteins detected are highlighted in blue, kinetochore proteins above the significance threshold are labelled. Components of the Set1b complex are highlighted in maroon and labelled. n = 4 biological replicates. (*c*) HeLa cells were fixed in paraformaldehyde and stained for Ndc80 (red), Bod1 (green), and a marker of the centromeric region (ACA, blue). A pair of kinetochores is shown. White line indicates the region chosen for a line profile (bottom panel) of fluorescence intensities across the kinetochore pair.

### Bod1 directly interacts with the Ndc80 complex

To confirm the Bod1-Ndc80 interaction detected by MS analysis, we performed pull down assays with purified Bod1-GST on Sepharose beads to validate Ndc80 as a bona fide Bod1 interactor. Ndc80 localises to kinetochores as part of the heterotetrameric Ndc80 complex, consisting of Ndc80, Nuf2, Spc24, and Spc25 [30,31] (figure 4*a*). Bod1-GST coated beads pulled out Ndc80, Nuf2 and Spc24 from mitotic HeLa cell lysates (figure 4*b*) (Spc25 was not tested). In order to determine whether this was a direct interaction with the complex, we tethered recombinant Ndc80 Bonsai, a truncated form of the complex containing a GST-Nuf2-Spc24 fusion and an Ndc80-Spc25 fusion that can be expressed in bacteria [32], to beads and incubated them with recombinantly expressed Bod1-MBP or MBP alone (figure 4*c,d*). Bod1-MBP interacted strongly with purified recombinant Ndc80 Bonsai complexes (figure 4*e*), supporting the proteomics and immunofluorescence data. Together these results demonstrate Bod1 is part of the outer kinetochore and directly interacts with Ndc80.

**Figure 4.**
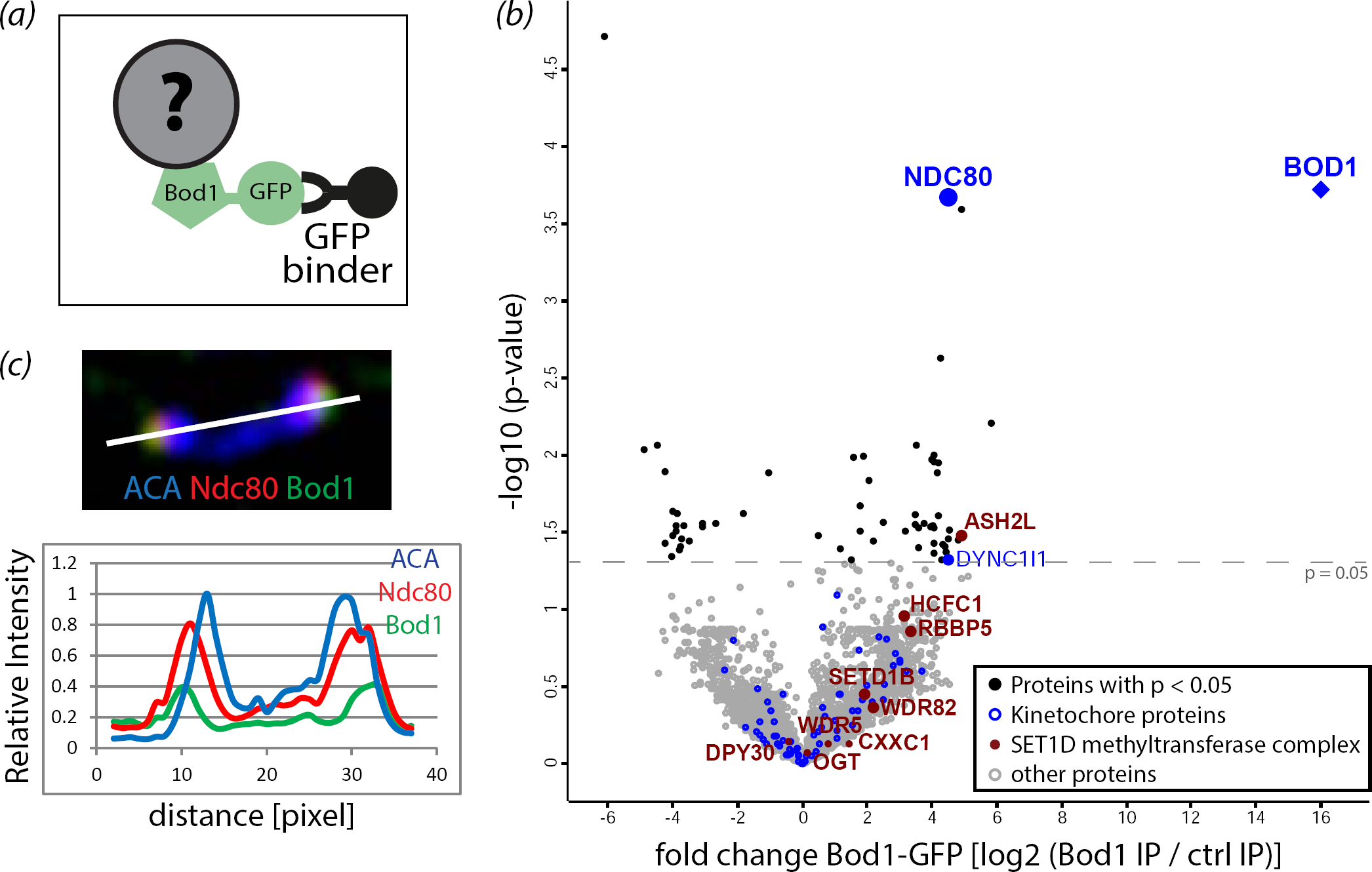
The Ndc80 complex interacts with Bod1 in mitotic HeLa cell extracts and *in vitro*. (*a*) Graphic representation of the full length Ndc80 complex. Regions of interest, including the microtubule and centromere binding regions of the complex, are labelled. (*b*) Co-precipitation of Ndc80 complex components Ndc80, Nuf2, and Spc24 from mitotic HeLa cell extracts with purified Bod1-GST. (*c*) Representation of the recombinant truncated Ndc80 Bonsai complex (as described in [32]). Ndc80-Spc25 and GST-Nuf2-Spc24 are expressed as fusion proteins. Both fusion constructs are co-expressed in *E. coli* from a dual-expression vector. (*d*) 150pmol recombinant Ndc80 Bonsai, consisting of dimers of one Ndc80-Spc25 fusion protein with one GST-Nuf2-Spc24 fusion protein, was immobilised on Sepharose beads and incubated with 1nmol Bod1-MBP or MBP. Binding was allowed for 1h. Proteins were resolved by SDS-PAGE and immunoblotted using simultaneous detection of the MBP (red) epitope tag on Bod1 and the GST (green) epitope tag on Nuf2-Spc24. (*e*) Amount of bound protein in (*d*) was quantified relative to the input. Statistical test used was Student’s t-test. Error bars represent standard error. n = 9 separate experiments.

### Ndc80 is essential for Bod1 kinetochore recruitment

To test if the Ndc80 complex was necessary for Bod1 kinetochore recruitment in cells, we depleted Ndc80 from HeLa cells using siRNA. Ndc80 depletion also reduced the immunofluorescence signal of its direct binding partner Nuf2 (figure 5). In contrast, we observed only a minor reduction in Knl1 signal, indicating that Ndc80 siRNA did not destabilise the entire outer kinetochore. Crucially, Ndc80 depletion resulted in significant loss of Bod1 from kinetochores, concomitant with an increase in localisation of B56. The increase in B56 localisation recapitulates our previous observations that siRNA depletion of Bod1 elevates B56 levels at kinetochores [9] and suggests that the localisation of Bod1 to kinetochores might limit PP2A-B56 accumulation at these sites.

**Figure 5.**
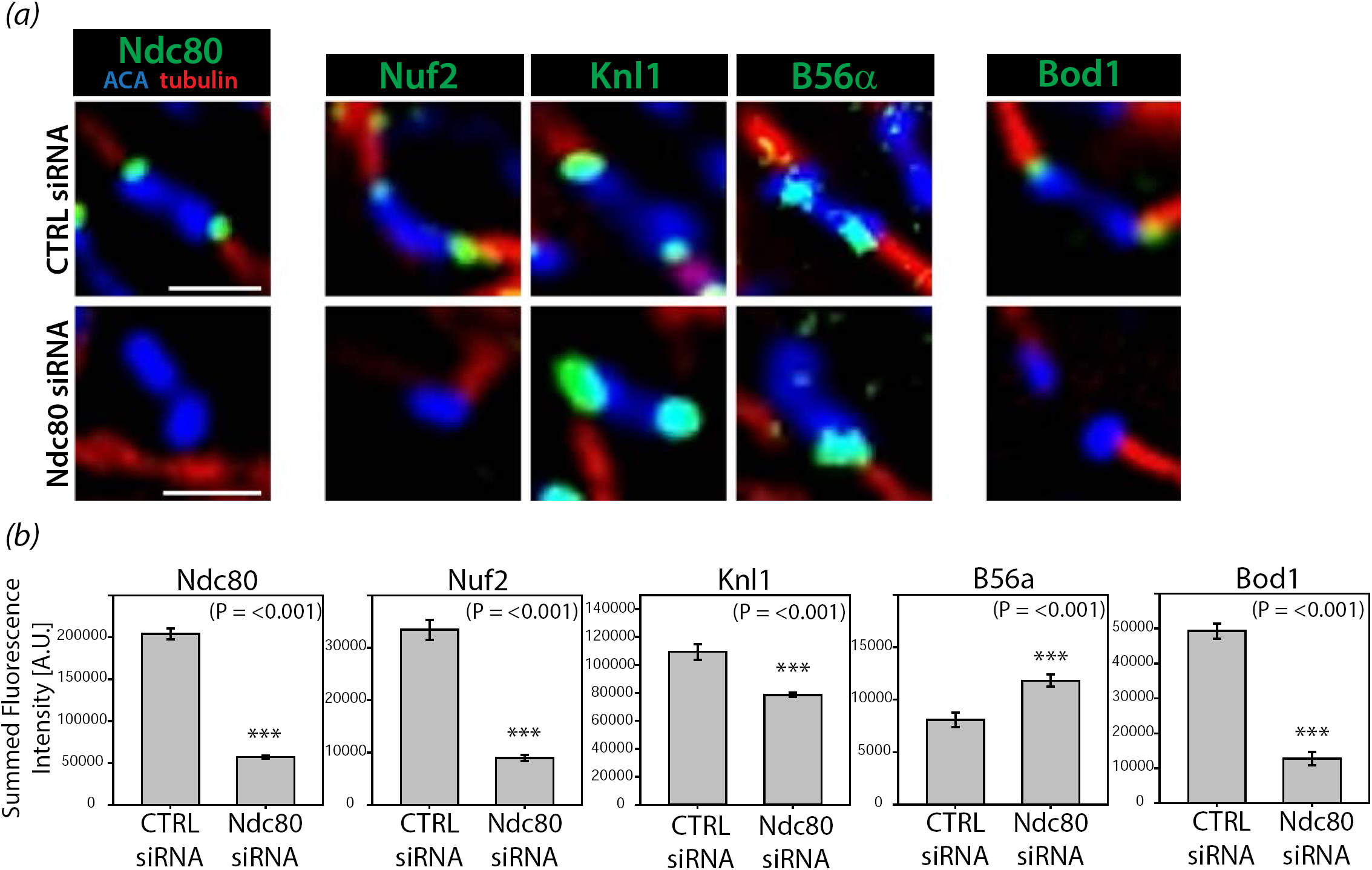
The Ndc80 complex is essential for Bod1 recruitment to mitotic kinetochores. (*a*) HeLa cells were treated with either control siRNA or Ndc80 siRNA. After 48h cells were fixed in paraformaldehyde and stained with the indicated kinetochore proteins (green). Kinetochores of chromosomes on the metaphase plate are shown. Localisation to the kinetochore, i.e. the interface between centromeric region (anti-centromere antibody, ACA, in blue) and spindle microtubules (tubulin, red) was compared. Scale bars are 1 μm. (*b*) Quantification of the images in (*a*). n = 10 cells per condition. Error bars represent standard error. Three asterisks indicate highly significant difference in Student’s t test.

### Bod1 depletion results in loss of Ndc80 phosphorylation at its N-terminal tail

siRNA-mediated depletion of Bod1 leads to mitotic arrest as cells are unable to maintain chromosome alignment in metaphase until anaphase onset [18]. This biorientation phenotype is accompanied by an increase in syntelic kinetochore-microtubule attachments, an attachment conformation in which a pair of sister kinetochores connects to spindle microtubules emanating from the same pole. This form of attachment can lead to erroneous mitosis, aneuploidy, and cell death [33] and therefore needs to be corrected before cells progress through mitosis. Correction of syntelic attachments is enabled by phosphorylation of outer kinetochore proteins [4,5,7,34], among them the N-terminus of Ndc80. Upon phosphorylation, the affinity of these proteins to microtubules is reduced and attachments are destabilised [5]. To evaluate if Ndc80 phosphorylation is dependent on Bod1 levels, we probed Ndc80 phosphorylation at its N-terminal serine 55 (S55) in Bod1 depleted cells using a phospho-specific antibody against this site. First, we compared control metaphase cells with Bod1 depleted cells that exhibited a clear biorientation phenotype. Bod1 depleted cells with biorientation phenotypes exhibited a 65% decrease in Ndc80 phosphorylation at S55 (figure 6*a,b*). However, Bod1 depletion and development of the bioriention phenotype also led to a small but significant reduction (∼30%) in the intensity of total Ndc80 protein at the kinetochore. We therefore aimed to understand the effects of Bod1 depletion on Ndc80 phosphorylation at S55 in cells just entering mitosis, before development of the mature chromosome misalignment phenotype.

**Figure 6.**
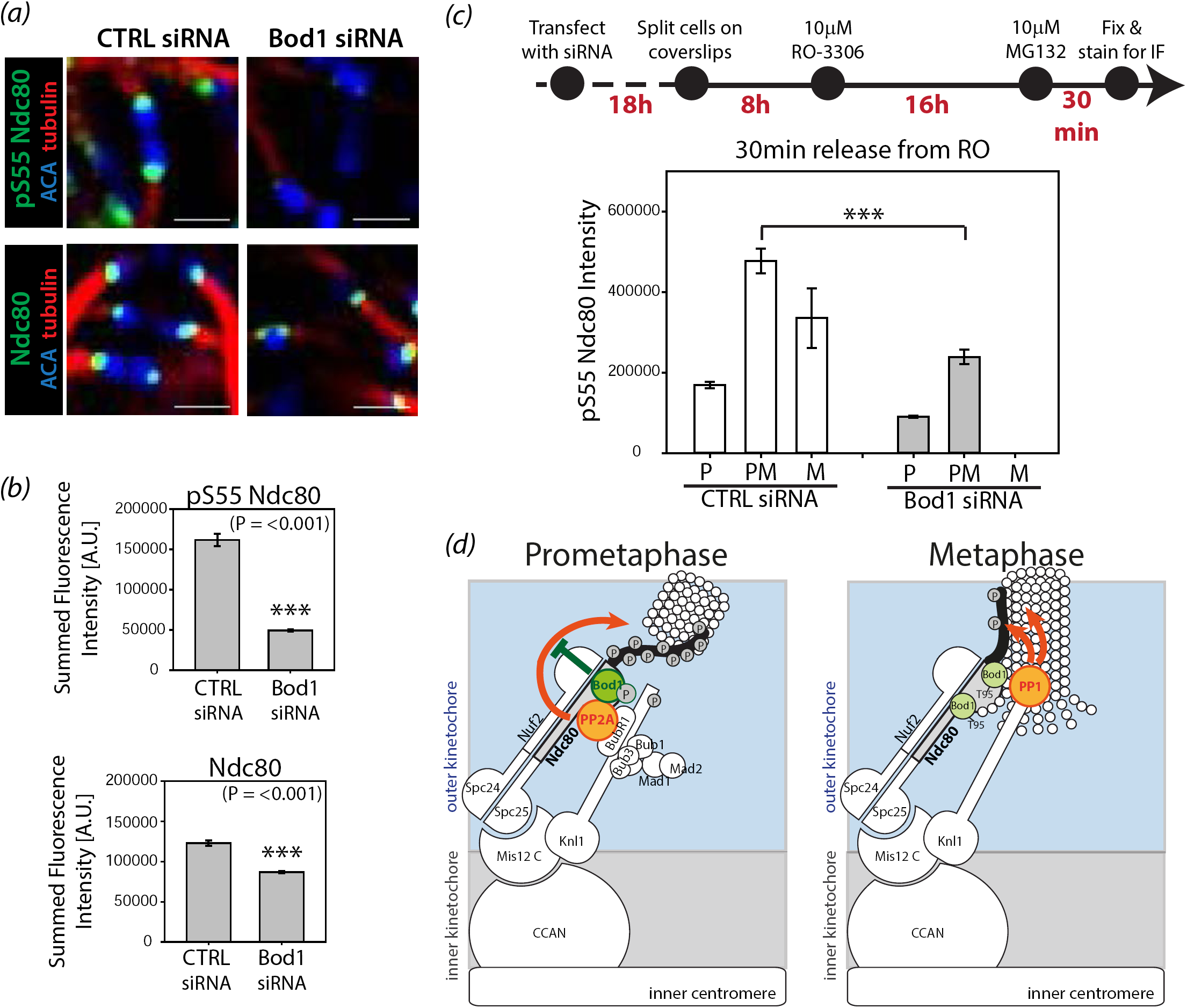
Bod1 depletion interferes with phosphorylation of the Ndc80 N-terminal tail. (*a*) HeLa cells were treated with control or Bod1 siRNA for 48h, fixed in paraformaldehyde and stained with a phospho-specific antibody against a phosphoepitope (pS55) in the Ndc80 N-terminal tail, or a total Ndc80 antibody. Metaphase cells or cells with the characteristic Bod1 chromosome misalignment phenotype were imaged. Kinetochores of chromosomes on the metaphase plate are shown. Scale bars are 1 μm. (*b*) pS55 and total Ndc80 intensities at kinetochores were quantified. n = 10 cells per condition. Statistical test used was Student’s t-test. Error bars represent standard error. Three asterisks indicate highly significant difference. (*c*) Timeline and quantification of treatments to compare kinetics of Ndc80 phosphorylation in control and Bod1-depleted HeLa cells in early mitosis. 26h after transfection, cells were synchronised at G2/M transition with the Cdk1 inhibitor RO-3306 for 16h. They were then released into culture medium containing the proteasome inhibitor MG132 to prevent mitotic exit and fixed 30min after their release from G1/M arrest. pS55 Ndc80 fluorescence intensities at kinetochores of mitotic cells were quantified. n = 50 mitotic cells per condition. Three asterisks indicate high significance in pairwise multiple comparison after ANOVA on ranks (p < 0.001). (*d*) A unified model integrating data presented in this paper with previous findings on the temporal regulation of kinetochore phosphatases in mitosis. Bod1 phosphorylation in early mitosis can inhibit PP2A-B56 activity [9] and prevent premature dephosphorylation of Ndc80 at its N-terminus. Dephosphorylation of Bod1 in later stages by an unknown mechanism, coincides with a surge in phosphatase activity, recruitment of PP1, and therefore enables PP1-mediated dephosphorylation of kinetochore epitopes in metaphase [12–14,49–51].

To selectively sample cells between nuclear envelope breakdown and metaphase, we first synchronised cells at the G2/M transition with RO-3306, a Cdk1 inhibitor [35]. The cells were then released into mitosis. Their medium was supplemented with the proteasome inhibitor MG132 to limit mitotic progression to prophase, prometaphase, and metaphase (figure 6*c*). Thirty minutes after RO-3306 release we assessed both Ndc80 pS55 kinetochore intensities and mitotic progression. At the early stages of mitosis, when error correction takes place, Bod1-depleted cells had significantly lower Ndc80 pS55 levels than control cells (figure 6*c*). Bod1-depleted cells also showed a delay in mitotic progression, even in these early stages of mitosis, which is consistent with previous findings [18].

Our results suggest a model where Bod1 holds PP2A-B56 activity in check during prometaphase. This ensures that phosphorylation of Ndc80 and possibly other kinetochore substrates can occur, allowing turnover of attachments (figure 6*d*). Once amphitelic kinetochore-microtubules attachment are achieved, we observe rapid dephosphorylation of Bod1 at T95. We have previously shown that loss of Bod1 phosphorylation prevents its inhibition of PP2A-B56 [9]. Active PP2A-B56 can then dephosphorylate the RVSF and SILK motifs on KNL1 [36], resulting in recruitment of PP1 [12] and stabilization of kinetochore-microtubule attachments by PP1-mediated dephosphorylation of Ndc80 [13].

## 3. Discussion

Using a combination of proteomics, *in vitro* recombinant interaction studies, and *in vivo* siRNA mediated localisation studies, we have demonstrated that the PP2A-B56 regulator Bod1 is recruited to kinetochores by Ndc80. Using quantitative immunofluorescence, we have also demonstrated that Bod1 is required to prevent premature dephosphorylation of Ndc80 during early mitosis. Together with MCAK S92 [15] and CENP-U/PBIP1 T78 [16], Ndc80 S55 adds to the list of phospho-epitopes at kinetochores lost upon Bod1 depletion. Bod1 therefore controls the phosphorylation of distinct groups of kinetochore proteins, all of which are implicated in the establishment of proper amphitelic kinetochore – microtubule attachments.

Bod1 is present at kinetochores throughout mitosis, from nuclear envelope breakdown until the end of anaphase [18]. Here, quantitative analysis has demonstrated that Bod1 localisation to kinetochores peaks during metaphase. Phosphorylation of Bod1 at T95 is essential for its inhibitory function against PP2A-B56 [9]. In contrast to the total population of Bod1, the pool of phopsho-T95 Bod1 peaked at kinetochores during prometaphase. This timing potentiates the inhibitory activity towards PP2A-B56 when phosphatase localisation to kinetochores is greatest and its activity needs to be properly regulated to allow correction of erroneous attachments [11].

While PP2A-B56 is recruited to kinetochores in a Knl1-dependent manner [16,17], we show here that Bod1 relies on the Ndc80 complex for kinetochore binding instead. Inter-complex interactions within the KMN network are an emerging theme in attachment regulation: The mitotic checkpoint kinase Mps1 is recruited to kinetochores through Ndc80 [14,37,38], but it phosphorylates MELT motifs on Knl1 [39–41]. Whether dynamic structural changes of the outer kinetochore allow interaction between components of the different KMN subcomplexes or medium affinity binding surfaces allow protein turn-over and localised gradients of diffusible proteins remains to be determined.

## 4. Materials and methods

### Cell lines and cell culture

HeLa S3 cells were maintained in EMEM (Lonza) and supplemented with 10% fetal calf serum, 2mM L-glutamine, 100U/ml penicillin and 100mg/ml streptomycin. A cell line stably expressing Bod1-GFP was generated using HeLa cells harbouring a single Flp recombination target (FRT) site in their genome (a kind gift from Patrick Meraldi [42]) and maintained in the media described above with an additional 200 mg/ml hygromycin. Cells lines were maintained at 37°C with 5% CO_2_ in a humidified incubator. For transfections, cells were seeded in 6-well dishes and transfected with 300ng plasmid DNA per well using Effectene transfection reagent (Qiagen) or with 33nM siRNA oligo duplexes or medium GC control siRNA (Invitrogen) using lipofectamine 2000 (Invitrogen). Cells were split onto coverslips the next day. Immunofluorescence staining of the cells and immunoblot analysis were performed 48h after siRNA transfection.

### Generation of peptide antibodies

Sheep were immunized with the immunogenic Bod1 peptide NH2-CRNGLRQ[pS]VVQS-COOH and serum containing the polyclonal antibody was collected in three batches. The third batch, obtained 91 days after the initial immunisation and 7 days after the third antigen booster injection, was used for antibody purification. 5mg NH2-CRNGLRQSVVQS-COOH peptide were added to 5ml activated Affigel-10 (Biorad) in 0.1M Na phosphate buffer pH7.8. Coupling of was allowed to take place over night. Then, residual iodoacetate groups were blocked with 0.2% β-mercaptoethanol and non-covalently bound peptide was removed by consequtive washes with 0.1M NaHCO3, 1M Na2CO3, water, 0.2M glycine-HCl pH2.0, 150mM NaCl, and TBS. The resin was stored in 0.1% NaN3 in TBS. For antibody purification, 4ml serum were diluted 1:1 with TBS and passed through a 0.2μm filter. The diluted serum was run over the column ten times. The column was then washed with TBS, 0.5M NaCl, 20mM Tris-HCl pH7.4, 0.2% Triton-X in TBS, and TBS. A low pH elution was performed with 0.15M NaCl, 0.2M glycine-HCl pH2.0 collecting 1ml fractions with each tube containing 0.1ml 2MTris-HCl pH8.5. After re-equilibrating the pH of the column by washing with TBS, a second, guadinuim hydrochloride elution was performed with 6M GuanidinHCl in TBS. Samples of all fractions were spotted onto nitrocellulose membranes and protein content was visualised with Ponceau S (Sigma). All fractions that contained antibody proteins were pooled and dialysed into TBS over night. Antibodies were stored in 0.1% sodium azide in TBS at 4°C.

### Immunofluorescence and microscopy

Cells were seeded on coverslips (thickness 1.5) 24h before fixation. Cells were pre-permeabilised with ice-cold cytoskeleton (CSK) buffer (100mM NaCl, 300mM sucrose, 3mM MgCl2, 10mM PIPES (pH6.8)) containing 0.1% Triton X-100 for 3min at 4°C before fixation with 37% paraformaldehyde in PBS at room temperature. Samples were re-hydrated with TBS-T before blocking with 1% normal donkey serum in AbDil (0.25% v/v Tween-20, 2% w/v BSA, 0.1% w/v NaN3 in TBS). Primary antibodies were added, diluted in AbDil, for 1h. Cells were carefully washed with TBS-T and secondary antibodies (1:500 in AbDil, Jackson ImmunoResearch) were added for 30min. Cells were washed again and 4',6-Diamidino-2-phenylindole (DAPI, Sigma) was added at 1μg/ml in TBS for 10min. Coverslips were washed with TBS and mounted onto microscope slides by inverting them into mounting medium (0.5 % p-phenylenediamine (Free Base; Sigma) in 20 mM Tris, pH 8.8, 90 % glycerol). Primary antibodies included polyclonal sheep Bod1 antibodies (0.5μg/ml), mouse anti-B56α (1:100, BD Biosciences), mouse anti-Ndc80 (1:500, Abcam [9G3]), mouse anti-Nuf2 (1:300, Abcam), rabbit anti-Dsn1 (1:300, GeneTex), rabbit anti-CASC5 [Knl1] (1:1000, Abcam), rabbit anti-Mlf1 [CENPU] (1:200, Rockland), rabbit anti-Ndc80 (phospho-Ser55) antibody (1:300, GeneTex), rat anti-tubulin (1:500, AbD Serotec), human anti-centromere autoantisera [ACA] (1:1000, a kind gift from Sara Marshall, Ninewells Hospital, Dundee). Secondary antibodies (Jackson ImmunoResearch) were used at 1:150. Three-dimensional deconvolution image data sets were acquired on a DeltaVision imaging system (Applied Precision) equipped with an Olympus 1-UB836 microscope, CCD camera (CoolSNAP_HQ/ICX285), and 100x/1.4 NA Plan-Apochromat oil immersion objectives (Olympus). Z stacks were collected 0.2μm apart and deconvolved using softWoRx (Applied Precision).

### Image analysis

Image data were imported into OMERO and quantification of kinetochore intensities was performed using OMERO.mtools [43]. Kinetochores were identified from deconvolved image stacks covering the whole cell by segmenting them based on anti-centromere antibody (ACA) staining using Otsu thresholding. The minimum object size was set to 50 pixels and the perimeter of the automatically generated mask was expanded by 4 pixels to include the outer kinetochore in the analysis. The fluorescence signal within this mask was measured and background staining was quantified in a 2 pixel annulus with a 1 pixel gap to the perimeter of each segmented mask. Fluorescence intensity at the kinetochore was then calculated as the summed fluorescence intensity within the mask subtracting the product of its size and the average background intensity in the 2 pixel annulus. Only positive values were taken into consideration for statistical analysis. All images were stored in OMERO, and figures were generated using OMERO.figure.

### Affinity purification and immunoblotting

For affinity purification, HeLa S3 cells were arrested in 5μM STLC for 18h. After gentle mitotic shake-off, cells were resuspended in lysis buffer (20mM Tris acetate pH 7.5, 1mM EGTA, 1mM EDTA, 10mM Na-β-glycerophosphate, 5mM Na-pyrophosphate, 1mM Na-orthovanadate, 50mM NaF, 1μM microcystin, 0.27M sucrose, 10μg/ml leupeptin, 10μg/ml pepstatin, 10μg/ml aprotinin), containing 0.01% and 0.05% Triton X-100 for stable and transient transfections respectively, and disrupted with four rounds of freeze fracturing. After depleting insoluble proteins by centrifugation (4°C, 10.000rpm, 5min), affinity purification was performed using GFP-Binder (Chromotek) for 90min at 4°C and constant agitation. Purified samples were washed and resolved by SDS-PAGE. Immunoblotting was performed using standard procedures, and secondary antibody was detected using either Clarity Western ECL Substrate (BioRad) and X-Ray films (Kodak) or the Odyssey Clx infrared detection system (LI-COR). Primary antibodies included mouse anti-B56α (1:500, Abcam), mouse anti-B56d (1:500, Abcam), mouse anti-Ndc80 (1:1000, Abcam [9G3]), mouse anti-Nuf2 (1:1000, Abcam), rabbit anti-Spc24 (1:1000, Abcam [EPR11548(B)]), mouse anti-MBP (1:20,000, NEB), goat anti-GST (1:5000, Abcam), mouse anti-Vinculin (1:10,000, Abcam [SPM227]), mouse anti-GFP (1:1000, Roche), rabbit anti-Bod1 (1:500, Abcam), polyclonal sheep anti-Bod1 (2ug/ml). Secondary antibodies were sheep anti-mouse IgG, HRP-linked (1:10,000, GE Healthcare), goat anti-rabbit IgG, HRP-linked (1:5000, Cell Signalling), donkey anti-goat IgG, HRP-linked (1:20,000, Promega), donkey anti-sheep HRP (1:20,000, Sigma), IRDye 680LT donkey anti-mouse IgG (H+L) (1:20,000, LI-COR), IRDye 800CW donkey anti-goat IgG (H+L) (1:20,000, LI-COR). LI-COR images were quantified using Image Studio software V2.0 (LI-COR), with signal intensity normalized to input protein levels.

### Mass spectrometry

Eight 15cm plates of stably Bod1-GFP or GFP transfected cells or two 15cm plates of transiently Bod1-GFP or GFP transfected cells were arrested in mitosis and affinity purification using GFP-Binder (Chromotek) was performed as described above. Proteins were eluted with 2xSDS buffer and the full eluate was run on a 4-12% SDS-PAGE. Bands were visualised using Coomassie Brilliant Blue and lanes were cut into 4 gel pieces. Gel pieces were subsequently de-stained with ammonium bicarbonate and acetonitrile as an organic solvent and dried completely in a vacuum centrifuge. 10mM DTT was used to reduce cystine disulphide bonds and the resulting thiol groups were irreversibly alkylated to S-carboxyamidomethylcysteine with 55mM iodoacetamide. Excess iodoacetamide was removed and gel pieces were dried in a vacuum centrifuge before enzymatic digest of the proteins. In-gel digest was performed with 20ng/μl trypsin in 50mM ammonium bicarbonate at 37°C o/n. Tryptic peptides were extracted from the gel by repeated addition of 0.1%TFA/acetonitrile extraction solution and sonication. Peptide samples were cleaned for mass spectrometry using a C18-Ziptip protocol. Mass spectrometry was performed on an LTQ Orbitrap Velos Pro instrument (Thermo Fisher Scientific). Mass spectrometry raw data were processed in the MaxQuant software package V 1.3.0.5 utilising the Uniprot Human database (09/08/2012) [44]. Parameters applied include: minimum peptide length = 7, Protein FDR = 0.01, Site FDR = 0.01. Peptides with variable modifications (N-terminal acetylation of the protein, oxMet, and pyroGlu) and fixed modifications (S-carboxyamidomethylcysteine) were accounted for in the analysis. Shotgun proteomics data analysis, including statistical analysis and GO term analysis, was performed using the Perseus software package V1.5.5.3 [45]. Statistical test performed was an unpaired Student’s t-test with a threshold p-value of 0.05.

### Protein expression and purification

Ndc80 Bonsai was expressed and purified as described previously [32]. For production of Bod1-MBP and MBP, 5ml LB medium containing the appropriate selection marker were inoculated with transformed BL21 *E. coli*. After 18h at 37°C, starter cultures were transferred into 2l conical flasks containing 500ml LB medium with the selection antibiotic. Cultures were grown in shaking incubators at 37°C up to OD600 = 0.4. After adding 100mM benzylalcohol for 30min at 37°C, recombinant protein production was induced by addition of 0.1mM Isopropyl β-D-1-thiogalactopyranoside (IPTG). Protein expression was allowed for 18h at 18°C. Bacteria were harvested by ultracentrifugation (5250xg, 4°C, 30min, slow deceleration)and lysed by resuspending them in PBS containing a protease inhibitor cocktail (Roche) and adding 1mg/ml lysozyme. Cells were incubated at 4°C under constant agitation for 30min after which Triton X-100 was added to a final concentration of 1%. The suspension was sonicated for 30s on ice and left to incubate another 30min at 4°C. The lysate was sonicated twice more and insoluble debris was pelleted by ultracentrifugation (26.000xg, 4°C, 1h). For protein purification, 1ml amylose resin (NEB) was pre-equilibrated with binding buffer (50mM Tris-HCl pH 7.5, 100mM NaCl, 1mM DTT) and the soluble fraction of the protein lysate was added after passing through a 0.2μm filter. Binding was allowed for 2h at 4°C under constant agitation. The recombinant protein bound to beads was washed with binding buffer. To elute the protein, 500μl binding buffer containing 20mM maltose were added and samples were incubated for 90min at 4°C under agitation. The supernatant was transferred into a Slide-A-Lyzer dialysis cassette (Pierce) and dialysed into interaction buffer (20mM Tris–HCl, 20mM NaCl, 10% glycerol, 1mM EGTA, 1mM DTT) over night. The concentration of dialysed protein was determined using a Bradford colorimetric assay. If protein concentrations were below 0.2mg/ml, protein solutions were concentrated using Vivaspin columns (GE Healthcare) at 4750rpm, 4°C. Proteins were aliquoted and stored at -80°C.

### Pull down experiments

150pmol Ndc80Bonsai, coupled to glutathione beads, were pre-incubated with 0.01% insulin in interaction buffer (20mM Tris–HCl, 20mM NaCl, 10% glycerol, 1mM EGTA, 1mM DTT, Complete protease inhibitors (Roche)) for 20min at 4°C. 1nmol MBP or Bod1-MBP was added to the beads and binding was allowed to take place for 1h at 4°C. After washing with interaction buffer, proteins were eluted with SDS loading buffer and all eluate was loaded for immunoblot analysis. 25pmols MBP or Bod1-MBP were loaded as input controls. Band intensity was determined using the ImageStudio software package. Total amount of protein in the pull down was determined by using the input as a reference.

### Statistical analysis

Statistical significance tests were performed using Sigma Plot 12.5 (Systat Software Inc.). For pairwise comparison, datasets were tested for normal distribution and then analysed by Student’s t test (for Gaussian distributions) or by Mann-Whitney U test (for Non-Gaussian distributions). For group-wise comparison, datasets were compared by Kruskal-Wallis one-way analysis of variance (ANOVA) on ranks, followed by pairwise multiple comparison procedures (Dunn's Method).

### siRNAs

Knl1 was depleted using 5′-GCAUGUAUCUCUUAAGGAA-3′ [36]. siRNA targeting Bod1 was 5’-GCCACAAAUAGAACGAGCAAUUCAU-3’ [18]. Ndc80 was depleted using an siRNA with the sequence 5’-AAGTTCAAAAGCTGGATGATCTT-3’ [46]. All isoforms of the B56 PP2A regulatory subunit were depleted using a pool of 5’-GCUCAAAGAUGCCACUUCA-3’ (B56α/PPP2R5A), 5’-CGCAUGAUCUCAGUGAAUA-3’ (B56β(PPP2R5B)), 5’-GGAUUUGCCUUACCACUAA-3’ (B56?/PPP2R5C), 5′-UCCAUGGACUGAUCUAUAA-3′ (B56δ/PPP2R5D), 5’-UUAAUGAACUGGUGGACUA-3’ (B56ε/PPP2R5E), described in [11]. Stealth^(tm)^ RNAi siRNA Negative Control, Med GC (Invitrogen) was used for control transfections.

## Data accessibility

The mass spectrometry proteomics data have been deposited to the ProteomeXchange Consortium [47] via the PRIDE partner repository [48] with the dataset identifier PXD006322.

## Acknowledgements

We are grateful to Jennifer G DeLuca for the gift of the Ndc80^Bonsai^ expression construct and Adrian T Saurin for the Knl1 siRNA. We thank the Dundee Centre for Advanced Scientific Technologies and Fingerprints Proteomics in Dundee for their support and all the members of our laboratory for helpful discussions.

## Author contributions

KS, IMP, and JRS conceived the experimental strategy. KS performed and analysed the experiments. StH performed and analysed the mass spectrometry with input from KS. KS, IMP and JRS wrote the manuscript.

## References

1. Schmid, M., Steinlein, C., Tian, Q., Hanlon Newell, A. E., Gessler, M., Olson, S. B., Rosenwald, A., Kneitz, B. & Fedorov, L. M. 2014 Mosaic variegated aneuploidy in mouse BubR1 deficient embryos and pregnancy loss in human. Chromosom. Res. 22, 375–392. (doi:10.1007/s10577-014-9432-x)

2. Holland, A. J. & Cleveland, D. W. 2012 Losing balance: the origin and impact of aneuploidy in cancer. EMBO Rep. 13, 501–14. (doi:10.1038/embor.2012.55).

3. Funabiki, H. & Wynne, D. J. 2013 Making an effective switch at the kinetochore by phosphorylation and dephosphorylation. Chromosoma. 122, 135–158. (doi:10.1007/s00412-013-0401-5)

4. Cheeseman, I. M., Anderson, S., Jwa, M., Green, E. M., Kang, J. seog, Yates, J. R., Chan, C. S. M., Drubin, D. G. & Barnes, G. 2002 Phospho-regulation of kinetochore-microtubule attachments by the Aurora kinase Ipl1p. Cell. 111, 163–172. (doi:10.1016/S0092-8674(02)00973-X)

5. Welburn, J. P. I., Vleugel, M., Liu, D., Yates, J. R., Lampson, M. A., Fukagawa, T. & Cheeseman, I. M. 2010 Aurora B Phosphorylates Spatially Distinct Targets to Differentially Regulate the Kinetochore-Microtubule Interface. Mol. Cell 38, 383–392. (doi:10.1016/j.molcel.2010.02.034)

6. Foley, E. a & Kapoor, T. M. 2013 Microtubule attachment and spindle assembly checkpoint signalling at the kinetochore. Nat. Rev. Mol. Cell Biol. 14, 25–37. (doi:10.1038/nrm3494)

7. Guimaraes, G. J., Dong, Y., McEwen, B. F. & DeLuca, J. G. 2008 Kinetochore-Microtubule Attachment Relies on the Disordered N-Terminal Tail Domain of Hec1. Curr. Biol. 18, 1778–1784. (doi:10.1016/j.cub.2008.08.012)

8. Vallardi, G. & Saurin, A. T. 2015 Mitotic kinases and phosphatases cooperate to shape the right response. Cell Cycle 14, 795–796. (doi:10.1080/15384101.2015.1006546)

9. Porter, I. M., Schleicher, K., Porter, M. & Swedlow, J. R. 2013 Bod1 regulates protein phosphatase 2A at mitotic kinetochores. Nat Commun 4, 2677. (doi:10.1038/ncomms3677)

10. Wurzenberger, C., Held, M., Lampson, M. A., Poser, I., Hyman, A. A. & Gerlich, D. W. 2012 Sds22 and repo-man stabilize chromosome segregation by counteracting Aurora B on anaphase kinetochores. J. Cell Biol. 198, 173–183. (doi:10.1083/jcb.201112112)

11. Foley, E. a., Maldonado, M. & Kapoor, T. M. 2011 Formation of stable attachments between kinetochores and microtubules depends on the B56-PP2A phosphatase. Nat. Cell Biol. 13, 1265–1271. (doi:10.1038/ncb2327)

12. Liu, D., Vleugel, M., Backer, C. B., Hori, T., Fukagawa, T., Cheeseman, I. M. & Lampson, M. A. 2010 Regulated targeting of protein phosphatase 1 to the outer kinetochore by KNL1 opposes Aurora B kinase. J. Cell Biol. 188, 809–820. (doi:10.1083/jcb.201001006)

13. Posch, M., Khoudoli, G. A., Swift, S., King, E. M., DeLuca, J. G. & Swedlow, J. R. 2010 Sds22 regulates aurora B activity and microtubule-kinetochore interactions at mitosis. J. Cell Biol. 191, 61–74. (doi:10.1083/jcb.200912046)

14. Nijenhuis, W. et al. 2013 A TPR domain-containing N-terminal module of MPS1 is required for its kinetochore localization by Aurora B. J. Cell Biol. 201, 217–231. (doi:10.1083/jcb.201210033)

15. Shi, Y. 2009 Serine/Threonine Phosphatases: Mechanism through Structure. Cell. 139, 468–484. (doi:10.1016/j.cell.2009.10.006)

16. Kruse, T., Zhang, G., Larsen, M. S. Y., Lischetti, T., Streicher, W., Kragh Nielsen, T., Bjørn, S. P. & Nilsson, J. 2013 Direct binding between BubR1 and B56-PP2A phosphatase complexes regulate mitotic progression. J. Cell Sci. 126, 1086–92.(doi:10.1242/jcs.122481)

17. Xu, P., Raetz, E. a, Kitagawa, M., Virshup, D. M. & Lee, S. H. 2013 BUBR1 recruits PP2A via the B56 family of targeting subunits to promote chromosome congression. Biol. Open 2, 479–86. (doi:10.1242/bio.20134051)

18. Porter, I. M., McClelland, S. E., Khoudoli, G. A., Hunter, C. J., Andersen, J. S., McAinsh, A. D., Blow, J. J. & Swedlow, J. R. 2007 Bod 1, a novel kinetochore protein required for chromosome biorientation. J. Cell Biol. 179, 187–197. (doi:10.1083/jcb.200704098)

19. Esmaeeli-Nieh, S. et al. 2016 BOD1 Is Required for Cognitive Function in Humans and Drosophila. PLoS Genet. 12. (doi:10.1371/journal.pgen.1006022)

20. Pan, D., Du, Y., Ren, Z., Chen, Y., Li, X. & Hu, B. 2016 Radiation induces premature chromatid separation via the miR-142-3p / Bod1 pathway in carcinoma cells. Oncotarget 7, 60432–60445. (doi:10.18632/oncotarget.11080)

21. Junttila, M. R. et al. 2007 CIP2A Inhibits PP2A in Human Malignancies. Cell 130, 51–62. (doi:10.1016/j.cell.2007.04.044)

22. Fan, L., Liu, M., Guo, M., Hu, C., Yan, Z., Chen, J., Chen, G. & Huang, Y. 2016 FAM122A, a new endogenous inhibitor of protein phosphatase 2A. Oncotarget 7, 63887–63900. (doi:10.18632/oncotarget.11698)

23. Li, M., Makkinje, A. & Damuni, Z. 1996 Molecular identification of I1PP2A, a novel potent heat-stable inhibitor protein of protein phosphatase 2A. Biochemistry 35, 6998–7002. (doi:10.1021/bi960581y)

24. Li, M., Makkinje, A. & Damuni, Z. 1996 The myeloid leukemia-associated protein SET is a potent inhibitor of protein phosphatase 2A. J. Biol. Chem. 271, 11059–11062. (doi:10.1074/jbc.271.19.11059)

25. Mcconnell, J., Gomez, R., Mccorvey, L., Law, B. & Wadzinski, B. 2007 Identification of a PP2A-interacting protein that functions as a negative regulator of phosphatase activity in the ATM/ATR signaling pathway. Oncogene 26, 6021–6030. (doi:10.1038/sj.onc.1210406)

26. Gharbi-Ayachi, A., Labbé, J.-C., Burgess, A., Vigneron, S., Strub, J.-M., Brioudes, E., Van-Dorsselaer, A., Castro, A. & Lorca, T. 2010 The substrate of Greatwall kinase, Arpp19, controls mitosis by inhibiting protein phosphatase 2A. Science 330, 1673–7. (doi:10.1126/science.1197048)

27. Mochida, S., Maslen, S. L., Skehel, M. & Hunt, T. 2010 Greatwall phosphorylates an inhibitor of protein phosphatase 2A that is essential for mitosis. Science 330, 1670–3. (doi:10.1126/science.1195689)

28. Suijkerbuijk, S. J. E. et al. 2012 The Vertebrate Mitotic Checkpoint Protein BUBR1 Is an Unusual Pseudokinase. Dev. Cell 22, 1321–1329. (doi:10.1016/j.devcel.2012.03.009)

29. van Nuland, R., Smits, A. H., Pallaki, P., Jansen, P. W. T. C., Vermeulen, M. & Timmers, H. T. M. 2013 Quantitative dissection and stoichiometry determination of the human SET1/MLL histone methyltransferase complexes. Mol. Cell. Biol. 33, 2067–77. (doi:10.1128/MCB.01742-12)

30. Bharadwaj, R., Qi, W. & Yu, H. 2004 Identification of Two Novel Components of the Human NDC80 Kinetochore Complex. J. Biol. Chem. 279, 13076–13085. (doi:10.1074/jbc.M310224200)

31. Ciferri, C. et al. 2005 Architecture of the human Ndc80-Hec1 complex, a critical constituent of the outer kinetochore. J. Biol. Chem. 280, 29088–29095. (doi:10.1074/jbc.M504070200)

32. Ciferri, C. et al. 2008 Implications for Kinetochore-Microtubule Attachment from the Structure of an Engineered Ndc80 Complex. Cell 133, 427–439.(doi:10.1016/j.cell.2008.03.020)

33. Gordon, D. J., Resio, B. & Pellman, D. 2012 Causes and consequences of aneuploidy in cancer. Nat. Rev. Genet. 13, 189–203. (doi:10.1038/nrg3123)

34. DeLuca, J. G., Gall, W. E., Ciferri, C., Cimini, D., Musacchio, A. & Salmon, E. D. 2006 Kinetochore Microtubule Dynamics and Attachment Stability Are Regulated by Hec1. Cell 127, 969–982. (doi:10.1016/j.cell.2006.09.047)

35. Vassilev, L. T., Tovar, C., Chen, S., Knezevic, D., Zhao, X., Sun, H., Heimbrook, D. C. & Chen, L. 2006 Selective small-molecule inhibitor reveals critical mitotic functions of human CDK1. Proc. Natl. Acad. Sci. U. S. A. 103, 10660–5. (doi:10.1073/pnas.0600447103)

36. Nijenhuis, W., Vallardi, G., Teixeira, A., Kops, G. J. P. L. & Saurin, A. T. 2014 Negative feedback at kinetochores underlies a responsive spindle checkpoint signal. Nature 16, 1257–1264. (doi:10.1038/ncb3065)

37. Saurin, A. T., van der Waal, M. S., Medema, R. H., Lens, S. M. A. & Kops, G. J. P. L. 2011 Aurora B potentiates Mps1 activation to ensure rapid checkpoint establishment at the onset of mitosis. Nat Commun 2, 316. (doi:10.1038/ncomms1319)

38. Zhu, T. et al. 2013 Phosphorylation of microtubule-binding protein hec1 by mitotic kinase aurora b specifies spindle checkpoint kinase mps1 signaling at the kinetochore. J. Biol. Chem. 288, 36149–36159. (doi:10.1074/jbc.M113.507970)

39. London, N., Ceto, S., Ranish, J. A. & Biggins, S. 2012 Phosphoregulation of Spc105 by Mps1 and PP1 regulates Bub1 localization to kinetochores. Curr. Biol. 22, 900–906. (doi:10.1016/j.cub.2012.03.052)

40. Shepperd, L. A., Meadows, J. C., Sochaj, A. M., Lancaster, T. C., Zou, J., Buttrick, G. J., Rappsilber, J., Hardwick, K. G. & Millar, J. B. A. 2012 Phosphodependent recruitment of Bub1 and Bub3 to Spc7/KNL1 by Mph1 kinase maintains the spindle checkpoint. Curr. Biol. 22, 891–899. (doi:10.1016/j.cub.2012.03.051)

41. Yamagishi, Y., Yang, C.-H., Tanno, Y. & Watanabe, Y. 2012 MPS1/Mph1 phosphorylates the kinetochore protein KNL1/Spc7 to recruit SAC components. Nat. Cell Biol. 14, 746–752. (doi:10.1038/ncb2515)

42. Klebig, C., Korinth, D. & Meraldi, P. 2009 Bub1 regulates chromosome segregation in a kinetochore-independent manner. J. Cell Biol. 185, 841–858. (doi:10.1083/jcb.200902128)

43. Allan, C. et al. 2012 OME Remote Objects (OMERO): a flexible, model-driven data management system for experimental biology. Nat. Methods 9, 245–253. (doi:10.1038/nmeth.1896)

44. Cox, J. & Mann, M. 2008 MaxQuant enables high peptide identification rates, individualized p.p.b.-range mass accuracies and proteome-wide protein quantification. Nat. Biotechnol. 26, 1367–72. (doi:10.1038/nbt.1511)

45. Tyanova, S., Temu, T., Sinitcyn, P., Carlson, A., Hein, M. Y., Geiger, T., Mann, M. & Cox, J. 2016 The Perseus computational platform for comprehensive analysis of (prote)omics data. Nat. Methods 13, 731–40. (doi:10.1038/nmeth.3901)

46. Martin-Lluesma, S., Stucke, V. M. & Nigg, E. A. 2002 Role of Hec1 in spindle checkpoint signaling and kinetochore recruitment of Mad1/Mad2. Science (80-.). 297, 2267–2270. (doi:10.1126/science.1075596)

47. Vizcaíno, J. et al. 2014 ProteomeXchange provides globally coordinated proteomics data submission and dissemination. Nat Biotech 32, 223–226. (doi:10.1038/nbt.2839)

48. Vizcaino, J. A. et al. 2016 2016 update of the PRIDE database and its related tools. Nucleic Acids Res. 44, D447–D456. (doi:10.1093/nar/gkv1145)

49. Trinkle-Mulcahy, L., Andersen, J., Yun, W. L., Moorhead, G., Mann, M. & Lamond, A. I. 2006 Repo-Man recruits PP1-gamma to chromatin and is essential for cell viability. J. Cell Biol. 172, 679–692. (doi:10.1083/jcb.200508154)

50. Rosenberg, J. S., Cross, F. R. & Funabiki, H. 2011 KNL1/Spc105 recruits PP1 to silence the spindle assembly checkpoint. Curr. Biol. 21, 942–947. (doi:10.1016/j.cub.2011.04.011)

51. Meadows, J. C., Shepperd, L. A., Vanoosthuyse, V., Lancaster, T. C., Sochaj, A. M., Buttrick, G. J., Hardwick, K. G. & Millar, J. B. A. 2011 Spindle checkpoint silencing requires association of PP1 to both Spc7 and kinesin-8 motors. Dev. Cell 20, 739–750. (doi:10.1016/j.devcel.2011.05.008)

